# Beyond residence time: quantifying factors that drive the spatially explicit filtration services of a “pristine” oyster population

**DOI:** 10.1101/2021.03.10.434772

**Authors:** M.W. Gray, D. Pinton, A. Canestrelli, N. Dix, P. Marcum, D. Kimbro, R. Grizzle

## Abstract

The Guana-Tolomato-Matanzas (GTM) system is a relatively pristine and well-flushed estuary in Northeastern Florida, USA and characterized as having an extraordinarily high abundance of oysters. Historically, dense populations of oysters, such as those found in GTM, are believed to play an important role in water filtration; however, few biofiltration studies have had access to such pristine populations. To quantify the filtration service *(FS)* of Eastern oysters *(Crassostrea virginica)* in GTM at several spatial scales (i.e. reef, watershed, estuary), we implemented a model that solves for the hydrodynamics and depletion of particulate matter passing over model oyster populations, the latter of which were derived from detailed bay-wide surveys. The model results suggested that oyster reefs populating the GTM play an important role in water quality by filtering ~60% of the estuary’s volume within its residence time. Our approach teases apart the role of reef size, residence time, particle concentration, and other physical factors on the generation of *FS* at different spatial scales. Downstream effects were found to be very important for estuary *FS*, which depend on the spatial distribution of the reefs in the GTM and local and estuarine-scale hydrodynamics. Therefore, the difference between “realized” *FS* and the “potential” *FS* of a given reef may be substantial when considering the complex hydrodynamic and connectivity among populations at several scales. Our model results provide clear and actionable information for management of these oyster populations and conservation of their ecosystem services.

## 1. Introduction

Oyster conservation and restoration is often motivated by the suite of ecosystem services thought to accompany robust populations. For example, oyster reefs are widely recognized as an important nursery ground for commercially and ecologically valuable species (Atlantic States Marine Fisheries Commission 2007; Coen et al. 2007; Coen and Humphries 2017). The filtration services *(FS)* that extend from the suspension-feeding activity of oysters are also highly sought after. As oysters feed, they remove suspended microparticulate material (~2 - 100 □m) from the water column (Newell and Langdon 1996), improving water quality and clarity. Additionally, the byproducts of their feeding activity (feces, pseudofeces, and urea) aids in benthic-pelagic coupling, nutrient cycling, and facilitates denitrification. Recognizing the numerous benefits of oyster *FS*, top-down control of primary production, and improved water quality is a frequently stated ecological goal of oyster restoration (Mann and Powell 2007), especially in eutrophic estuaries and bays (Cranford 2019). Due to the substantial investment required for large-scale restoration or long-term conservation (Hernández et al. 2018), ecosystem models have become an increasingly popular tool to predict the ecological outcomes prior to any efforts.

Several notable ecosystem models have been developed over the past few decades to describe the role of oysters in controlling primary production. As models achieve greater sophistication, there has been greater emphasis to use the more ecologically realistic values for how oyster reefs interact with the overlying environment during their parameterization. It is worthwhile to note how the ecological modeling community has evolved while also acknowledging some remaining deficits. One important ecophysiological trait to account for during model creation is the role of environmental conditions on oyster filtration activity. Many laboratory studies have demonstrated oysters express elevated filtration rates under optimal laboratory conditions. Early modeling attempts used these elevated feeding rates (e.g., Newell 1988; Gerritsen et al. 1994), but subsequently have been criticized for their lack of ecological accuracy (Pomeroy et al. 2006; Mann and Powell 2007; Pomeroy et al. 2007; Cranford et al. 2011). Oysters living in the dynamic conditions found in estuaries often feed at slower and at more variable rates over time than those found in many laboratory studies (Grizzle et al. 2008; Cranford et al. 2011; M. W. Gray and Langdon 2018); thus, *in situ*-based feeding rates are considered more appropriate when modeling the effects of large populations on water quality. Furthermore, there are few examples of water filtration data that extend from fully-mature reefs because most native populations are functionally extinct (Beck et al. 2011), and even the stated goals for “restored” populations are far less dense (e.g. Allen et al. 2011) than the enormous and “pristine” populations described in early accounts by Euro-American settlers (Kurlansky 2007) or models reconstructing their demographics (Mann et al. 2009).

Aside from biological constraints on oyster *FS*, it is critically important to account for and incorporate hydrodynamics during model creation. Many previous biofiltration models have simplified the hydrodynamics and assumed these systems to be well-mixed and homogenous. However, accounting for mixing, heterogeneous water flow over reefs, and refiltration of water by oysters over time allows for a more precise estimate of time that oysters have to remove suspended material from the water column (Pomeroy et al. 2006; Fulford et al. 2007). Improved estimates of water exposure to oysters can lead to substantially different estimates of *FS* provided by oyster reefs. For example, Gray et al. (2019) estimated native Olympia oysters to filter 28% of Yaquina Bay, OR within a single residence time after accounting for hydrodynamics. This estimate is substantially larger than that of an earlier study (1% per residence time) by zu Ermgassen et al. (2013) who used a much simpler method when accounting for hydrodynamics (tidal prism method), which likely underestimated the residence time of the ecosystem (Lemagie and Lerczak 2015). Aside from residence time, the frequency at which a parcel of water was exposed to filter-feeding activity of oysters before exiting the estuary, termed encounter rate by Gray et al. (2019), was also considered to be important when estimating oyster *FS* but was not quantified. Water that repeatedly encounters oysters increases opportunity for refiltration by downstream reefs, but this effect can only be accounted for after knowing the precise location of oyster reefs and hydrodynamics.

The approach one uses to estimate spatially explicit oyster *FS* can also have a direct impact on the resulting estimates. All things being equal, larger populations will filter greater quantities of water than smaller ones, which does not provide much insight on the quality and relative services provided by subpopulations. Accounting for the area of populations when estimating *FS* enables one to determine which populations/locations are more efficient at removing seston. Furthermore, since clearance rates are non-linearly driven by the size of animals (i.e. dry tissue weight; DTW) and bound to be affected by density, reefs of similar area can have vastly different *FS* if they differ in terms of demographics. More often than not, detailed surveys of populations (especially historic ones) are lacking and demographic information is course, so assumptions about animal size and reef density during model formulation is derived from generalized relationships found in the literature (e.g. Mann et al. 2009; zu Ermgassen et al. 2012; zu Ermgassen et al. 2013). Accounting for the patchiness common among oyster reefs and demographics can help resolve ecosystem-scale *FS* and identify populations/locations that are more efficient at particle removal. Such information would greatly aid resource managers prioritizing reefs for conservation and/or developing restoration strategies that maximize return on *FS* after investment.

The historical role of oysters in exerting top-down control over primary productivity remains ambiguous and more resolved models are needed to understand oyster biofiltration at ecosystem scales. In this study, we sought to explore the filtration services of oysters in Guana-Tolomato-Matanzas River Estuary (GTM hereafter) in Northeastern Florida, USA. A model was created by exploiting recent advances in both biomonitoring and hydrodynamic modeling in the GTM. The GTM is home to an expansive population of Eastern oysters, *Crassostrea virginica*. In fact, high resolution surveys of reef boundaries and reef demographics have determined subpopulations to be very dense (1855 individuals m^-2^). Furthermore, the overall coverage of oysters within the intertidal and subtidal portion of the GTM estuary is small (4% of wet area), but due to the high density of animals found in reefs, the average density of oysters across the area of the estuary (50.7 oysters m^-2^) is among the higher estimates of historical populations (1880-1910) across the Atlantic Coast (range: 1.5 - 57.5 individuals m^-2^; zu Ermgassen et al. 2012), possibly resembling a “pristine” population that is capable of providing pre-colonial levels of *FS* (Mann et al. 2009).

## 2. Methods

### 2.1. Study Site

The GTM National Estuarine Research Reserve (GTMNERR) spans 60 km north and south of the city of St. Augustine in Northeastern Florida (Figure 1), at the transition between subtropical and temperate climates. The GTM estuary is primarily fed from the Atlantic Ocean through the St. Augustine inlet (29°91’N, 81°29’W) and Matanzas inlet (29°71’N, 81°23’W). It is traversed northsouth by the Intracoastal Waterway (ICW) through the Matanzas and Tolomato Rivers. The absence of major freshwater rivers makes the estuary well mixed and well flushed (Sheng et al. 2008). The three largest tributaries are Pellicer Creek, which empties into the Matanzas River in the southern portion of the estuary, San Sebastian River, which flows through the city of St. Augustine and empties into the Matanzas River, and Guana River, the northern reaches of which were impounded in the mid-1950s. Other minor tributaries are the Moultrie Creek and Moses Creek, which empty into the Matanzas River ~9 and ~17 km south of St. Augustine. The average tidal range in the estuary is ~1.5 m (NERRS 2021). Salinity varies from near zero ppt in the tributaries to 25-35 ppt near the inlets (NERRS 2021). Water temperature typically ranges from 15 to 30 °C (NERRS 2021). Dominant habitats in the estuary include salt marshes, mangroves, intertidal oyster reefs, tidal creeks, mudflats, and open water (Dix et al. 2017; Bacopoulos et al. 2019; Dix et al. 2019). Intertidal habitats are protected from ocean energy by barrier islands and dune systems.

**Figure 1.**
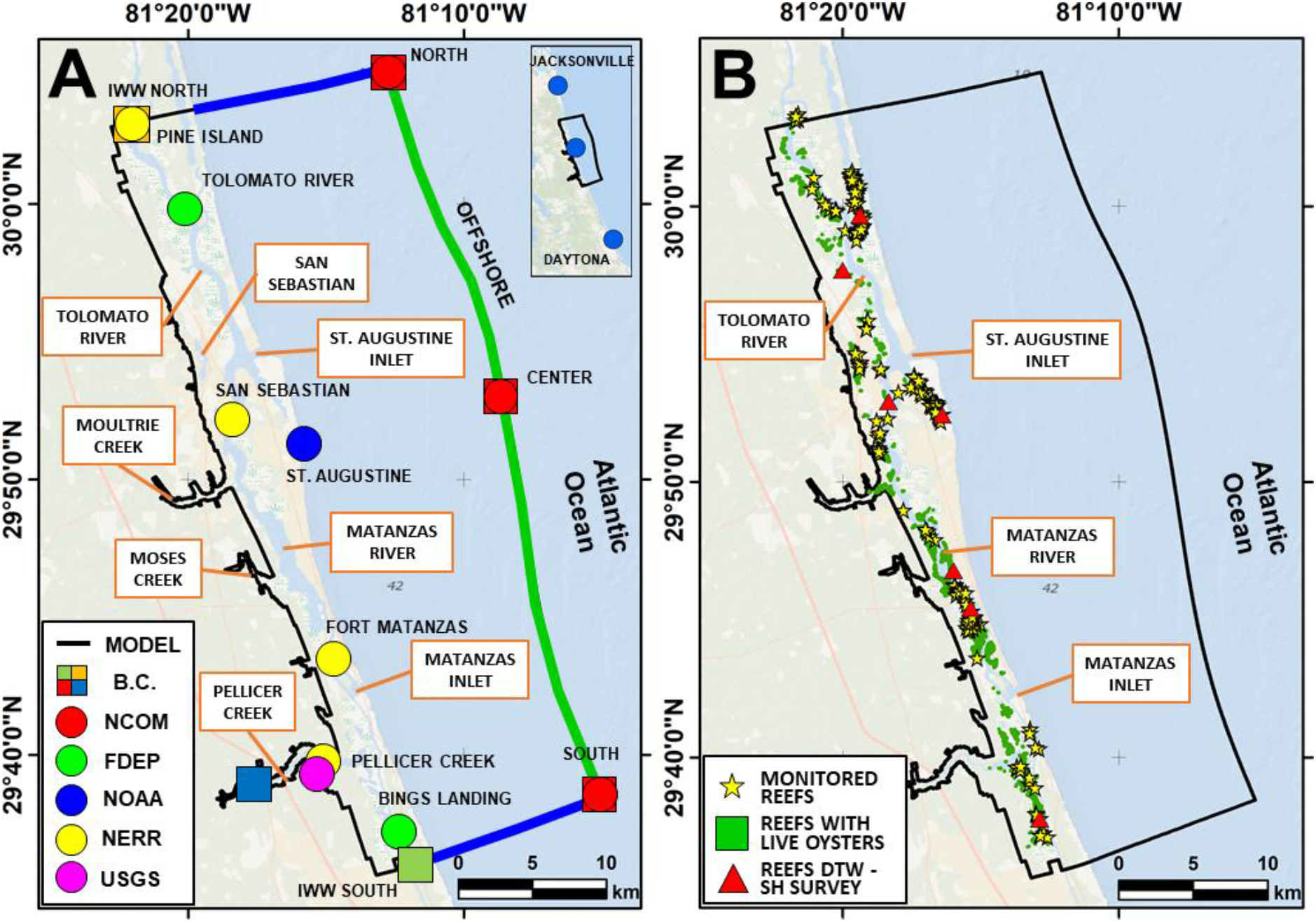
(A) The geographic position of the data sources (stations and numerical model points) used to determine the boundary conditions for our simulations (dots), and the geographic position of the open boundaries of the model domain (squares). The red dots indicate the locations where we extracted the boundary conditions for the water temperature. The other dots indicate the FDEP, NOAA, and NERR stations where we extracted the hydrodynamic boundary conditions. (B) Spatial distribution of the oyster reefs in the GTM estuary. Green areas indicate the reefs extracted from the Fish and Wildlife Research Institute (FWRI) database (https://hub.arcgis.com/datasets/myfwc::oyster-beds-in-florida), which are populated by live oysters. The yellows stars indicate the reefs surveyed by the GTMNERR. The red triangles indicate the reefs where we surveyed DTW and SH, to determine a relationship between them. In all plots, the black line represents the model domain.

### 2.2. Hydrodynamic model details

We solved for the hydrodynamics in the GTM estuary by using the Delft3D-FLOW model (https://oss.deltares.nl/web/delft3d/download). Delft3D-FLOW solves the Navier-Stokes equations for an incompressible fluid under the shallow water assumption and the Boussinesq approximation. It calculates non-steady flow resulting from the tidal and meteorological forcing on a regular, boundary-fitted grid. The latter allows for the accurate description of water level, currents, and transported solutes in topographically complex areas, such as the salt marshes of the GTM estuary. The model can handle wetting and drying of the grid cells due to tidal fluctuations. Delft3D-FLOW also computes water temperature using a heat flux model. The model calculates the heat exchange through the free water surface, and it includes the effects of solar radiation, convection, evapotranspiration, and precipitation.

In this study, we used a structured curvilinear grid that covers an area of ~1050 km^2^. The model domain is shown in Figure 1 (black line). The domain envelops the GTMNERR and was centered in the city of St. Augustine, FL, USA. The numerical grid describes: (i) the GTM estuary, composed of the ICW (Tolomato River and Matanzas River) and the Guana River up to the Guana Dam; (ii) the principal and minor affluents of the GTM in the study area (Pellicer Creek, Moultrie Creek, Moses Creek, and San Sebastian River); (iii) the Atlantic Ocean, up to ~12 km from the coastline, where the ocean bottom reaches ~20 m below MSL; (iv) the inlets of St. Augustine and Matanzas. The grid resolution was not uniform in the study domain; it was more refined along the estuary and inlets and less refined in the ocean and marshes. The average grid cell dimension varied from ~30 m × 100 m in the ocean to ~15 m × 20 m in the estuary.

The model bathymetry for the ocean was based on the National Oceanic and Atmospheric Administration (NOAA) data. The bathymetry for the GTM was based on the Florida Natural Areas Inventory (FNAI) vegetation map (https://www.fnai.org/LandCover.cfm), the United States Geological Survey (USGS) bathy LiDARs, the Army Corps topo-bathy LiDARs, and the NOAA LiDAR datasets (https://coast.noaa.gov/dataviewer/#/).

The model used hydrodynamic and water quality boundary conditions. At the offshore boundary (green lines in Figure 1A), we applied the harmonic constituents of the astronomical tide. The constituents were measured at three local NOAA stations placed along the coastline (see Appendix A - blue dots in Figure 1A). At this boundary, we also applied the water temperature extrapolated from the Regional Navy Coastal Ocean Model (NCOM-red dots in Figure 1A). Both boundary conditions were prescribed at three support points, indicated by the red squares (Figure 1A), which divide the boundary into two segments (green lines in Figure 1A). Points that lie in between each couple of support points were calculated by linear interpolation of the forcing at both ends. At the southern boundary of the ICW (green square in Figure 1A), we applied the water level and the water temperature measured by the Florida Department of Environmental Protection (FDEP) at the “Bing’s Landing” station (green dot in Figure 1A). At the northern boundary of the ICW (orange square in Figure 1A), we applied a Neumann boundary condition for the water level. Here, the alongshore water level gradient was assumed to be zero. At this boundary, we also applied the water temperature measured by the GTMNERR at the “Pine Island” station (yellow dot in Figure 1A). For Pellicer Creek (blue square in Figure 1A), we applied the tidally filtered discharge rate from the local USGS station (magenta dot in Figure 1A). Finally, we applied the meteorological forcings, corresponding to relative humidity, air temperature, wind direction, wind speed, precipitation, and solar radiation, to the entire domain. These data were measured by the GTMNERR meteorological station “Pellicer Creek” (yellow dot in Figure 1A).

For this study, we simulated a period of 30 days, which contained ~2 neap and ~2 spring tides (see Appendix A). The simulated period lasted from May 9^th^, 2018 to June 10^th^, 2018. The simulation time step was one minute.

To calculate the distribution of the residence time and the *FS* in the estuary, we interpolated the model statistics obtained for the simulated period on a uniform 50 m×50 m grid. The statistics we considered were the mean, minimum, and maximum water depth and the depth-averaged water temperature. Using the water depth, we identified the cells that are flooded at least once in the simulated period.

### 2.3. Oyster reefs

#### 2.3.1. Field surveys and allometric functions

We used the Fish and Wildlife Research Institute (FWRI) database (https://hub.arcgis.com/datasets/myfwc::oyster-beds-in-florida) to identify the geographic properties of the oyster reefs in the GTM estuary. Clipped to the study area boundary, the database contained ~4300 reefs divided into two classes: alive and dead. Dead reefs were distinguished as exposed mounds of disarticulated, bleached white shells mostly along the ICW channel (Garvis et al. 2020). In this study, we considered only the live reefs (Figure 1B). Detailed surveys were conducted between 2014 and 2020 by the GTMNERR to measure oyster population metrics (i.e., shell height and oyster density) over a sample of ~240 reefs (yellow stars in Figure 1B). The survey methods are described in Marcum et al. (2018). In short, oysters were collected from 2–3 0.25 m × 0.25 m (0.0625-m^2^) quadrats on each reef. Quadrats were placed randomly on 6–30-m transects (depending on reef size) laid along the elevation of the reef that appeared densest. This approach was used, based on the objective of the GTMNERR oyster survey, to minimize within-reef variability and the level of sampling effort required to detect regional patterns. Although transect placement was based on perceived live oyster density, transects curved with the shape of the reef and included both dense and less dense areas. Once collected, oysters were rinsed and clusters were broken apart to ensure all live oysters were counted. Shell height was measured from the umbo to the distal end of the largest shell with calipers on either a subset of 50 oysters or all oysters in each sample. Although the survey approach used by GTMNERR raises a potential bias for oversampling healthy portions of each reef (relative to total area of oyster polygon); the GTMNERR survey approach was used consistently across the estuary. Consequently, the relative differences between actual oyster reef densities and their estimates should be similar across sites.

Using the oyster dataset, we calculated the average oyster density *(D_Oys_*) and shell height *(SH)* for each surveyed reef. These parameters correspond to the number of animals per reef square meter and the average length of their shell in millimeters. We used ArcGIS to calculate their values on the not-surveyed reefs by using an inverse distance weighted (IDW) interpolation method. IDW predicts the values for the unsurveyed reefs by using the surrounding surveyed locations.

Filtration rates were dependent on the average dry tissue weight *(DTW)* of oysters in a given reef. Mean *DTW* were derived from relationships between *DTW* and *SH* from surveys conducted at seven stations distributed throughout the estuary (Figure 1B). Specifically, in June 2018, we haphazardly sampled three reefs separated by at least 10 m within each station (21 reefs total), yielding three oysters within ten different *SH* size classes (0–10, 11–20, 21–30, 31–40, 41–50, 51–60, 61–70, 71–80, 81–90, 91–100 mm) at each station. Oysters were cleaned of all epifauna, frozen, and then transported to Northeastern University for processing: oyster *SH* was determined by measuring the length (mm) of the longest bottom valve axis from ubmo to tip; *DTW* was quantified by shucking oysters, separating tissue from shell, placing tissue tin pre-weighed tin (Metler-Toledo Balance, model MS403S), drying the container at 60 °C for 72 hours, re-weighing the tin container, and subtracting pre- and post-dried container weight (g).

Non-linear regression analysis was used to determine that slope estimates between *DTW* and *SH* were similar among sites, indicating that a general relationship across estuary was permissible. Using Akaike Information Criterion (Akaike 1973) during non-linear model selection, the following three parameter exponential relationship between *DTW* and *SH* was found best to fit the data (R^2^ = 0.87):

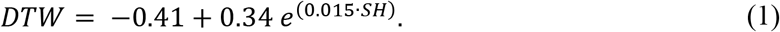

We then estimated the *DTW* in grams of the average oyster populating each reef using the local average *SH* as determined through surveys and applied it to Equation (1).

#### 2.3.2. Physiology

Oyster filtration rate *(FR_o_*) was defined as the volume of seawater filtered per unit time by each animal. The method was based on the approach proposed by zu Ermgassen et al. (2013) to examine the present and historical services of individual oysters along the Atlantic and Gulf Coasts. In their approach, *FR_o_* was estimated as:

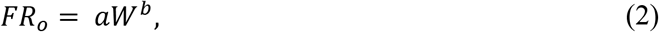

where *a* is the maximum filtration rate of an individual, and *b* is a scaling exponent. *b* describes how filtration scales with the dry tissue weight of animals (*W*, in grams), calculated from the individual shell height using the allometric function proposed by Newell and Langdon (1996). After careful analysis, zu Ermgassen et al. (2013) set *a* to 8.02 and *b* to 0.58. The latter is the universal value for suspension-feeding bivalves (Cranford et al. 2011). To account for the effect of temperature on the oyster, Equation (2) was modified using the method proposed by Cerco and Noel (2005) to:

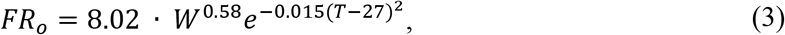

where *T* is the water temperature in Celsius degrees.

We then calculated the filtration rate of the average oyster populating each of the ~4300 reefs in the GTM estuary (Section 3.3.1) by applying the *DTW* calculated from Equation (1) to Equation (3).

To calculate the number *N* of animals populating a reef, we multiplied the local oyster density *(D_Oys_)* for the reef area (*A_Reef_*). We then calculated the filtration rate of the entire reef *(FR)* by multiplying the number of oysters populating it by the filtration rate of a singular animal *(FR_o_*).

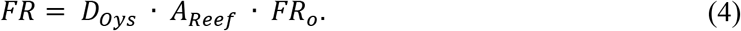

*FR_o_* is defined as the volume of seawater filtered per unit time per oyster (“o”, m^3^s^-1^oyster^-1^). *FR* is defined as the seawater volume filtered per unit time by an entire reef (m^3^s^-1^).

### 2.4. Residence Time calculation

To calculate the residence time in the study area, we employed a Lagrangian approach. In particular, we tracked the motion of virtual particles released in the GTM estuary by using the PART module of Delft3D. To simulate the motion of the particles, Delft3D-PART uses the hydrodynamic fields calculated by the FLOW module. This study employed conservative and neutrally buoyant particles, which were distributed uniformly in the GTM estuary. Particles were injected six times in the estuary, with a time interval of two hours between two consecutive injections. This method was used to cover the first tidal cycle and to consider the effect of tidal variability in the motion of the particles. The injection locations were the midpoints of the 50 m × 50 m regular grid cells, flooded for at least a time step of the hydrodynamic simulation. The output of the particle tracking module was a data file with the location of each particle within the estuary at each time step. The time step we chose for particle tracking was one minute, consistent with the hydrodynamic model.

In this study, we calculated three residence times: (i) the local residence time *(RT_L_*), defined for each 50 m × 50 m cell in the estuary, (ii) the watershed residence time *(RT_W_*), calculated for the nine watersheds we identified in the GTM estuary from the FDEP Waterbody ID drainage basin layer (https://geodata.dep.state.fl.us/datasets/waterbody-ids-wbids) and, (iii) the estuary residence time *(RT_E_*). The watersheds were identified by aggregating the ~40 watersheds located in the study domain, in nine groups (Figure 2A). The groups contain the afferent area of the most important rivers and creeks of the GTM and of the two inlets. In particular, the watersheds contain the afferent area of: the Tolomato River (W1), the Guana River (W2), the San Sebastian River (W3), the St. Augustine inlet (W4), the Salt Run (W5), the Moultrie Creek with the northern part of the Matanzas River (W6), the Moses Creek with the central part of the Matanzas River, above the tidal node (W7), the Matanzas inlet with the central part of the Matanzas River, below the tidal node (W8), and the Pellicer Creek with the southern part of the Matanzas River (W9). To calculate the local residence time, we identified all the particles entering each 50 m × 50 m cell, and the total time they spent inside the cell throughout the entire simulation. For each cell, the average of these times was the local residence time. The watershed and the estuary residence times were defined as the time needed for the particles to decrease their number by 1/e (with e ≈ 2.7) in the watersheds and estuary, respectively. These residence times were computed by considering only the particles released with the first injection.

**Figure 2.**
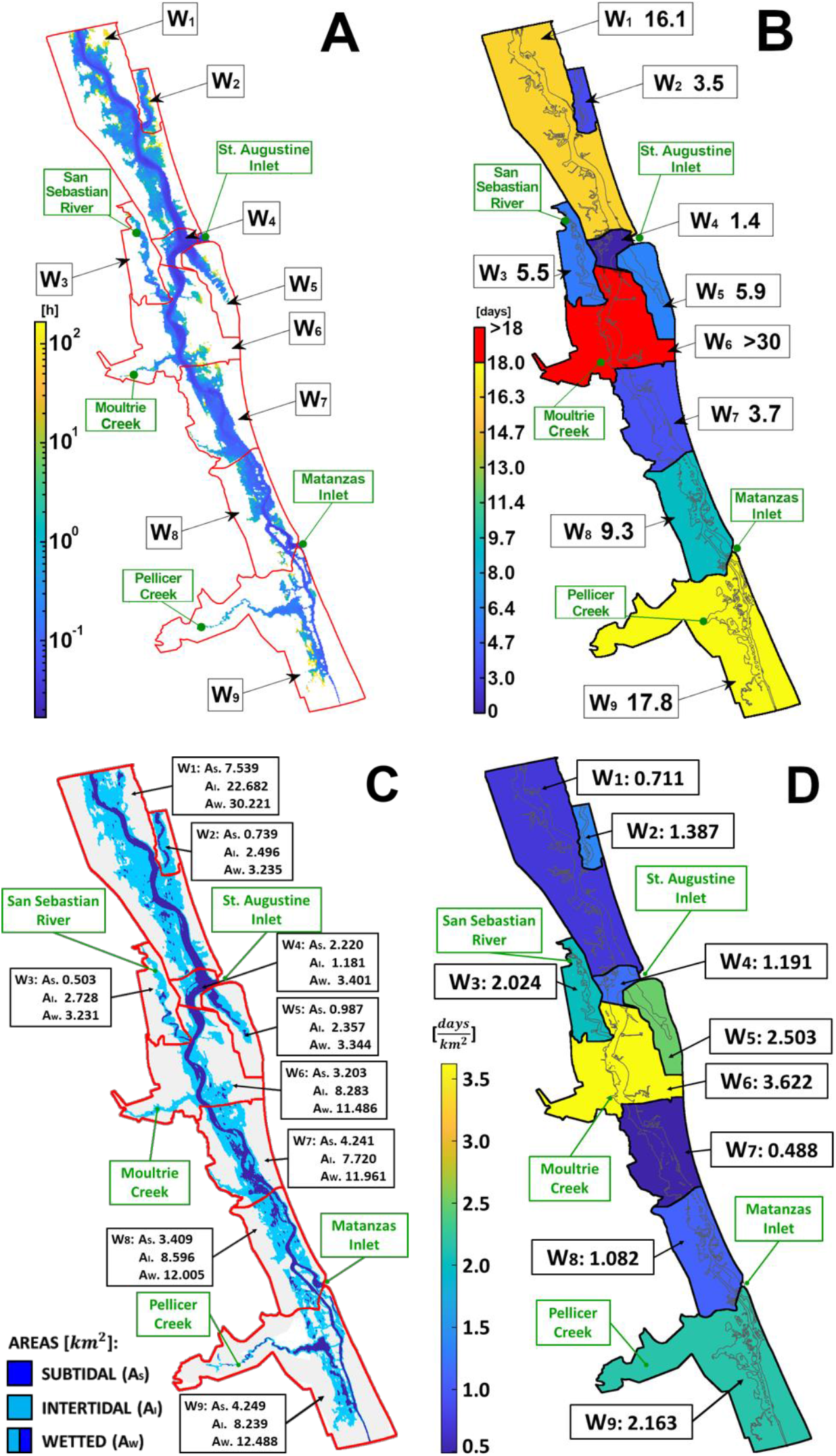
(A) Distribution of the local residence time *(RT_L_*) in the GTM estuary, using a 50 m × 50 m regular grid. Times are indicated in hours. The nine watersheds in which we divided the GTM are indicated in red. Note that the residence time is computed for the portion of the watershed that is wet over a spring-neap cycle. (B) Distribution of the watershed-scale residence times *(RT_W_*) calculated for the most important watersheds (Wi, i=1,…9) constituting the GTM. Times are indicated in days. (C) Distribution of the subtidal (-d_s_), intertidal (*A_I_*) and wetted (AW, or total) areas calculated for the most important watersheds (Wi, i=1,…9) constituting the GTM. (D) Distribution of the watershed-scale residence time per unit of intertidal watershed 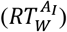, calculated for the most important watersheds (Wi, i=1,…9) constituting the GTM. Residence time is indicated in days/km^2^.

### 2.5. Filtration Services calculation

For a specific oyster population, *FS* were defined as the percentage of water mass filtered in the estuary within a single residence time. *FS* was computed at the levels of a single reef *(FS_R_*), an entire watershed *(FS_W_*), and at the estuary scale *(FS_E_*). To quantify the contribution of each reef to the estuary-scale *FS,* we developed a MATLAB code that processes the particles tracked by Delft3D-PART. The method was an improved version of the one proposed in Gray et al. (2019). Their code was based on the following assumptions: (i) each particle is initialized with a particle concentration of 1, (ii) at each time step, the concentration of the suspended particles is reduced by oyster reefs proportionally to a filtration rate (named “clearance rate” in Gray et al., 2019), (iii) there is no increase in the concentration of particles above the initial concentration.

The main difference between our approach and that of Gray et al. (2019) was conditions around assumption (ii). Gray et al. (2019) filtered particles at the cell scale. They divided their study area (Yaquina Bay, OR, USA) in a regular 150 m × 150 m horizontal grid. The area covered by reefs is usually a percentage of the cell area. This way, oysters filter particles even if they do not directly travel over the reef. Correction factors are thus needed to compute the correct filtration rates. The factors were estimated in Gray et al. (2019) as the proportion of the cell area occupied by the oysters. In reality, these factors depend on the local hydrodynamics. In our code, we overcame this issue and improved model resolution by estimating filtration at the reef spatial scale. We consider this a more realistic approach since we filtered only particles that travel over the polygon describing the reef. Moreover, Gray et al. (2019) does not consider the spatial variability of the oyster population properties (shell height) or oceanographic features (temperature) to calculate the filtration rate at the population level due to lack of data availability. Due to the information available to us and described above, we calculated *FR* using Equation (4) while accounting for the spatial distribution of water temperature, oyster density, and oyster dry tissue weight in the GTM estuary.

The amount of material *(dx)* removed by a reef due to oyster filtration was described with the following equation:

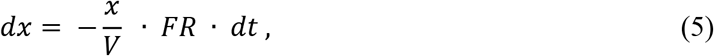

where the total amount of material in the area of interest of a reef is *x*, the volume of water above a reef is *V,* and the total filtration rate provided by the reef is *FR. FR* was calculated as described in Section 2.3.2. *dt* is the time step of the hydrodynamic and particle tracking simulations (one minute). The water volume on a reef (F) varied at each time step and depended on both the reef elevation and the water level calculated in the cells. Thus, knowing the reef properties and the water depth at any given time step, it was possible to calculate from Equations (4) and (5) the fractional change *(F_j,i_*) in the particle mass over any reef *j,* at any given time step *i.* Given the mass *x_i_* of the *I^th^* particle at the beginning of the time step, and knowing that the particle is suspended over the *j^th^* reef for that time step, the mass at the beginning of the next time step is:

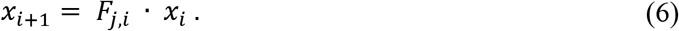

The MATLAB code records the amount of particle mass cleared by each reef at each time step. This allowed us to compute the total amount of particle mass removed from the estuary by each reef and to identify the reefs that most contribute to the filtration of the GTM estuary. The proportion of the estuary cleared by the *j^th^* reef *(FS_R,j_*) is:

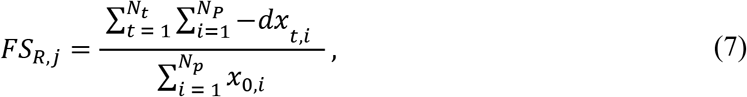

where *N_t_* is the total number of time steps (*t*) of the particle tracking simulation, and *N_p_* is the total number of particles injected in the estuary.

This definition of *FS* accounts for downstream effects because the *FS* of each reef depends on the filtration history of each particle. Because of the complex hydrodynamics of the estuary, due to the massive presence of salt marshes (Bacopoulos et al., 2019), and the dominant effect of the tide on the water fluxes (Sheng et al., 2008), the distribution of the downstream effect in the estuary is non-uniform.

### 2.6. Statistical analysis

#### 2.6.1. Genetic Algorithm

To evaluate the effect of the reef properties on their contribution to the estuary-scale *FS*, we performed a statistical analysis using a Genetic Algorithm (GA) (Madár et al. 2005). The GA simulates a biological evolution process. The process started with a population of random individuals, which grew at each time step until they reached an optimal solution. The individuals of each generational step were chosen using a fitness function calibrated on a target population. The optimal solution was achieved when significant changes in the individuals constituting the successive generations were negligible. In this study, the individuals were calculated using the following predictors, calculated for the particles entering the oyster reefs: (i) the average volume filtration rate of the reef per unit of reef area 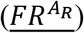; (ii) the average concentration (**<u>C</u>**) of the particle entering the reef. The concentrations were calculated as 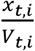; (iii) the average time spent by the particles on the reefs, which is the reef-scale residence time *(RT_R_*); (iv) the number of particles entering the reefs (*N*). Predictors (i) and (ii) were calculated averaging the values obtained from Delft3D and from the MATLAB algorithm, only at the first entry of the particles in the oyster reefs. The parameters (or model predictors) were used in the GA, to determine which of their combinations better described the target population. The changes in the population over the generations were the changes in the linear regression function used by the algorithm to fit the input data; the fitness function was the root mean square error (RMSE). Finally, the *FS* per unit of reef area 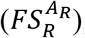, constitute the target population of the GA. They were obtained for each reef from *FS_R_,* which was calculated as described in Section 3.5, as follows:

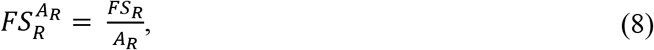

where *A_R_* is the area of a reef.

## 3. Results

### 3.1. Filtration rates

Physiological rates and other biological traits varied among oysters populations in each subestuary (Table 1). Estimates of oyster filtration rates ranged from 2.18 to 3.74 l hr^-1^. Averaged across the estuary oysters were estimated to clear 2.5 l hr^-1^ (SD: 0.54). After adjusting clearance rates to weight-standardized filtration rates of the small animals (average shell height 35 mm, average *DTW* estimate 0.17 g) that dominated reefs, we estimated that small animals clear on average 13.4 l h^-1^ g^-1^ to l7.5 1 h^-1^ g^-1^ across the estuary. Average weight-standardized filtration rates were fairly similar across all subestuaries with values that ranged 15.9 l h^-1^ g^-1^ (SD: 1.56).

**Table 1.**
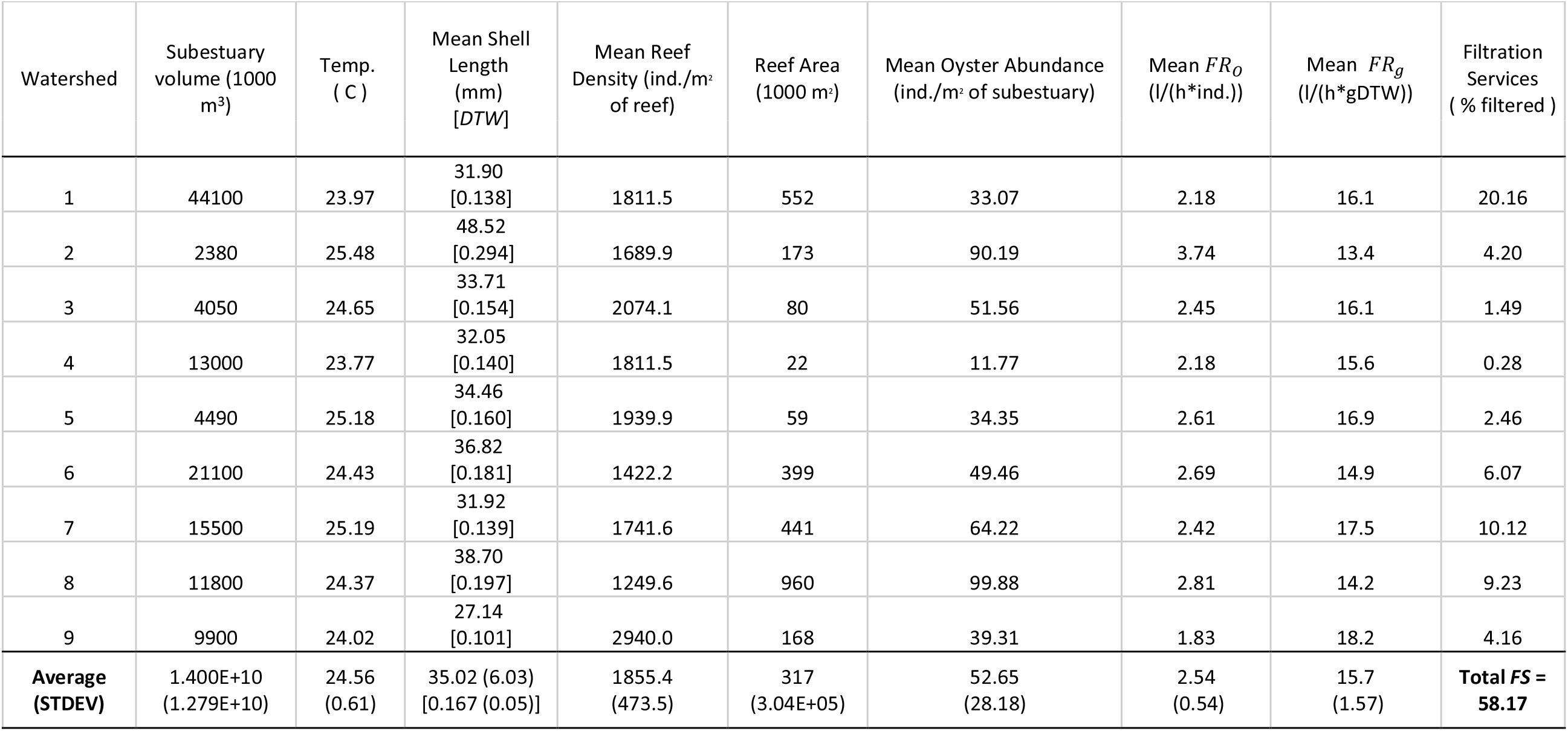
Watershed, subestuary, and subpopulation characteristics within the Guana-Tolomato-Matanzas (GTM estuary. Filtration services represent the percent of the GTM estuary that is filtered in by each subestuary. Mean Dry tissue weight (DTW) is in grams.

| Watershed | Subestuary<br>volume (1000<br>m<sup>3</sup>) | Temp.<br>( C ) | Mean Shell<br>Length<br>(mm)<br>[DTW] | Mean Reef<br>Density (ind./m<sup>2</sup><br>of reef) | Reef Area<br>(1000 m<sup>2</sup>) | Mean Oyster Abundance<br>(ind./m<sup>2</sup> of subestuary) | Mean $FR_o$<br>(l/(h*ind.)) | Mean $FR_g$<br>(l/(h*gDTW)) | Filtration<br>Services<br>(% filtered ) |
| --- | --- | --- | --- | --- | --- | --- | --- | --- | --- |
| 1 | 44100 | 23.97 | 31.90<br>[0.138] | 1811.5 | 552 | 33.07 | 2.18 | 16.1 | 20.16 |
| 2 | 2380 | 25.48 | 48.52<br>[0.294] | 1689.9 | 173 | 90.19 | 3.74 | 13.4 | 4.20 |
| 3 | 4050 | 24.65 | 33.71<br>[0.154] | 2074.1 | 80 | 51.56 | 2.45 | 16.1 | 1.49 |
| 4 | 13000 | 23.77 | 32.05<br>[0.140] | 1811.5 | 22 | 11.77 | 2.18 | 15.6 | 0.28 |
| 5 | 4490 | 25.18 | 34.46<br>[0.160] | 1939.9 | 59 | 34.35 | 2.61 | 16.9 | 2.46 |
| 6 | 21100 | 24.43 | 36.82<br>[0.181] | 1422.2 | 399 | 49.46 | 2.69 | 14.9 | 6.07 |
| 7 | 15500 | 25.19 | 31.92<br>[0.139] | 1741.6 | 441 | 64.22 | 2.42 | 17.5 | 10.12 |
| 8 | 11800 | 24.37 | 38.70<br>[0.197] | 1249.6 | 960 | 99.88 | 2.81 | 14.2 | 9.23 |
| 9 | 9900 | 24.02 | 27.14<br>[0.101] | 2940.0 | 168 | 39.31 | 1.83 | 18.2 | 4.16 |
| **Average**<br>**(STDEV)** | **1.400E+10**<br>**(1.279E+10)** | **24.56**<br>**(0.61)** | **35.02 (6.03)**<br>**[0.167 (0.05)]** | **1855.4**<br>**(473.5)** | **317**<br>**(3.04E+05)** | **52.65**<br>**(28.18)** | **2.54**<br>**(0.54)** | **15.7**<br>**(1.57)** | **Total FS =**<br>**58.17** |

### 3.2. Local residence time

Figure 2A shows the spatial distribution of the local residence time *(RT_L_*) on the GTM estuary. The figure shows that the Guana, Tolomato, and Matanzas River had the lowest residence times, ranging between 1 and 7 minutes. The lowest *RT_L_* values, ranging between 1 and 2 minutes, were observed next to St. Augustine and Matanzas inlets. These values gradually increased in the major watercourses and reached their maxima at the salt marshes, where the water velocity was lower than in the main channels. In these areas, the residence time went from ~30-240 minutes (0.5-4 hours) at the marsh platform to ~2-10 minutes at the marsh edge, where it was reduced due to the water exchange with tidal flats and channels flanking the marsh. The highest residence times were computed for the marshes farther from the inlets and adjacent to the mainland. In this area, *RT_L_* reached values up to ~5000-8000 minutes (~3.5-5.5 days). Compared to the northern part of the domain, we observed relatively higher values of *RT_L_* in the southern part of the GTM estuary. These low values were due to the smaller cross-section of the Matanzas River, in comparison with the Tolomato; the larger extension of salt marshes in comparison with the northern part of the estuary; and, the shallow depths at Matanzas inlet, which reduce tidal exchange with the sea.

### 3.3. Watershed scale residence time

The watershed-scale residence times *(RT_W_)* were calculated on the major watersheds of the GTM, described in Section 3.4, and are shown in Figure 2B. The different values of *RT_W_* were related to the temporal variation of the particle number in the watersheds, shown in Figure 3 as a percentage of their initial number in each watershed. *RT_W_* attained its lower value, equal to 1.4 days, for watershed W4, due to its proximity to St. Augustine inlet (Figure 2B). A much larger value was computed for the watershed W8, containing Matanzas inlet. Here the residence time was equal to 9.3 days. This difference was due to the lower fluxes moving through the shallow Matanzas inlet compared to St. Augustine inlet, the higher presence of salt marshes in the Matanzas watershed, and the greater area of watershed W8 in comparison with watershed W4. The larger oscillations observed in Figure 3F for watershed W4 indicated stronger tidal dominance in this watershed, with respect to W8 (Figure 3H). Higher residence times were obtained for the watersheds W3 and W5, which contain the San Sebastian River and Salt Run. These watersheds had a more peripheral location and lower water velocity than the major channels, which were connected to the inlets. The *RT_W_* were equal to 5.5 and 5.9 days, respectively. However, the small extension of salt marshes and the proximity to the Matanzas inlet and St. Augustine inlet kept the *RT_W_* values relatively small. Similar values were obtained for watersheds W2 and W7, where *RT_W_* was equal to 3.5 and 3.7 days, respectively.

**Figure 3.**
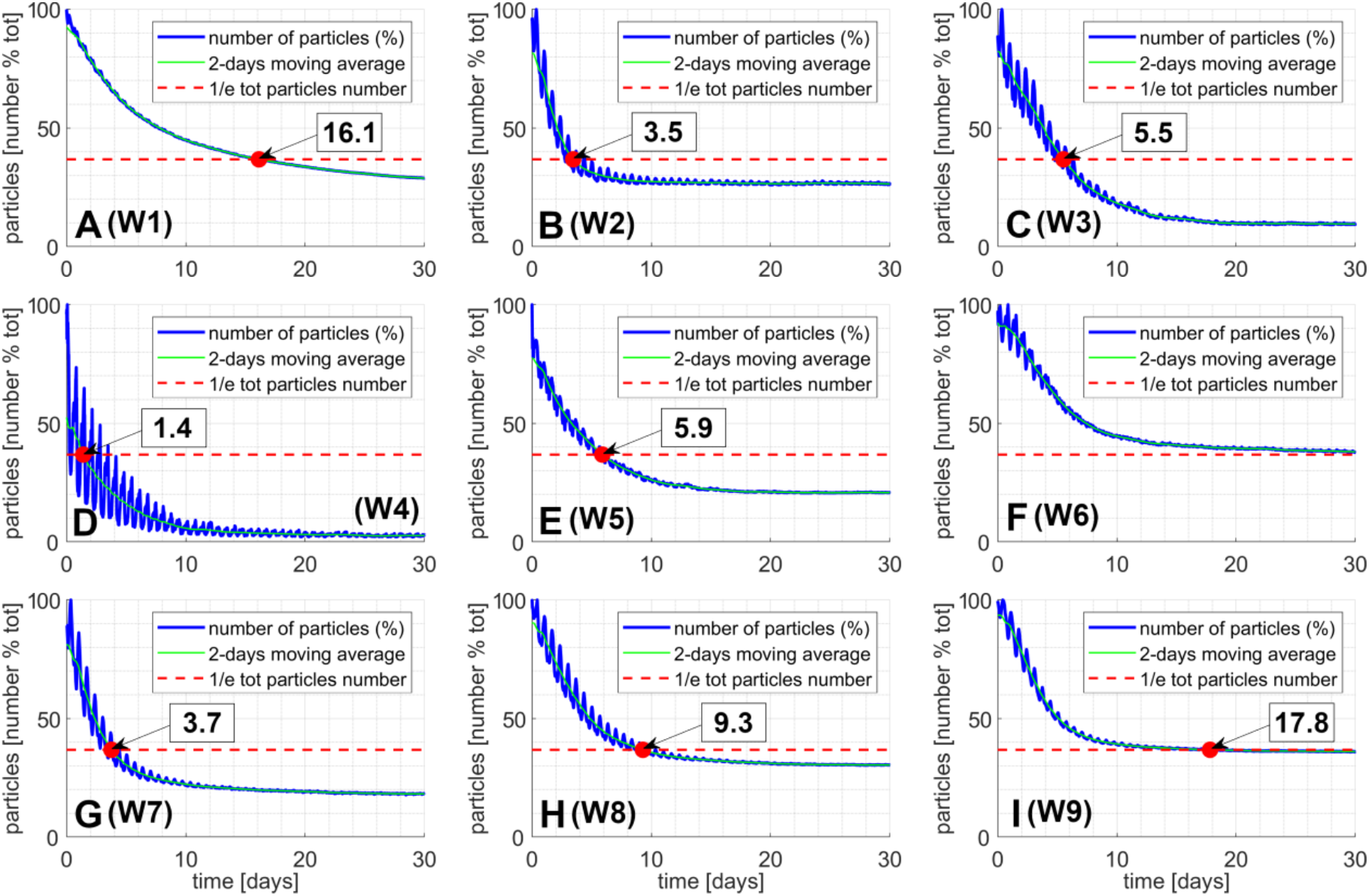
Each plot shows the number of particles located in each of the nine major watersheds divided by their initial number as a function of time. The red circle indicates when the number of particles reaches 1/e of the initial value, which is the watershed-scale residence time *(RT_W_*). The white boxes contain the value of the residence time expressed in days.

Residence times were larger for the watersheds located further away from the two inlets. Watershed W1, closer to St. Augustine inlet, had a residence time of 16.1 days, shorter than watershed W9, which was controlled by Matanzas inlet and had a residence time of 17.8 days. Watershed W6 did not reach the 1/e concentration of the initial number of particles in the 30-days simulation (Figure 3F). This was due to the massive presence of salt marshes, whose vegetation reduces the flow velocity, and Moultrie Creek, one of larger tidal tributaries.

It is important to note that, given the flow velocity, the residence time increased with the spatial dimension of the basin. To make *RT_W_* independent from the basin dimension, we calculated 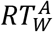, which was the watershed-scale residence time per unit of wetted watershed (*A_w_*). Hereinafter, “wetted” values of 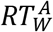 indicate the part of the watershed that was underwater at any instant of a neap-spring cycle. The wetted area of a watershed is composed of a subtidal portion (*A_s_*), which is always under the water level, and an intertidal portion *(A_I_*), which is flooded only for high water levels. The distribution of *A_W_*, *A_S_* and *A_I_*, in the GTM is shown in Figure 2C. Figure 2D shows the values of *RT_W_* per unit of intertidal area 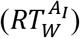 for the watersheds constituting the GTM. Similarly to Figure 2B, the distribution of 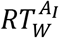 is non-uniform. A low 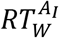 was observed for W4, equal to 1.191 days/km^2^, due to the proximity to St. Augustine inlet. The lowest value, equal to 0.488 days/km^2^, was observed for watershed W7 due to the significant extension of its wetted watershed, mostly composed of salt marshes, and to the low value of *RT_W_*. For the same reason, a low 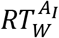, equal to 0.711 days/km^2^, was observed also for watershed W1. This was in contrast with the high value of *RT_W_* for W1 (the second highest in the GTM). The significant extension of the wetted watershed coupled with the proximity to Matanzas inlet, was the reason for the low 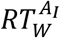 (1.082 days/km^2^) observed for W8. A similar 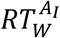, equal to 1.387 days/km^2^, was calculated for W2, containing the Guana River. This was due to the influence of the Tolomato River, the fluxes of which boosted the transport of particles from the Guana. Intermediate values of 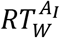, equal to 2.024, 2.503 and 2.163 days/km^2^, were observed for the watersheds W3, W5 and W9, which included Sebastian River, Salt Run, and Pellicer Creek, respectively. For W3 and W5, the limited extension of the intertidal watershed produced 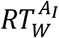 values higher than the ones observed in the previously described watersheds. For W9, the high 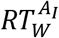 is due to the high *RT_W_*, which is the highest in the GTM. Finally, the highest value of 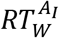 was observed for watershed W6. The value of 3.622 days/km^2^ for W6 was due to the high *RT_W_*, which is linked to the large presence of lateral intertidal regions, where the particles remain stuck after the flood phases.

### 3.4. Estuary residence time

Figure 4 shows the temporal variation of the particle numbers in the GTM estuary, calculated as a percentage of their initial number. From the numerical simulation, the estuary residence time *(RT_E_*) was estimated to be 12.6 days. This value was similar to the residence time observed by Sheng et al. (2008) in the region (~14 days).

**Figure 4.**
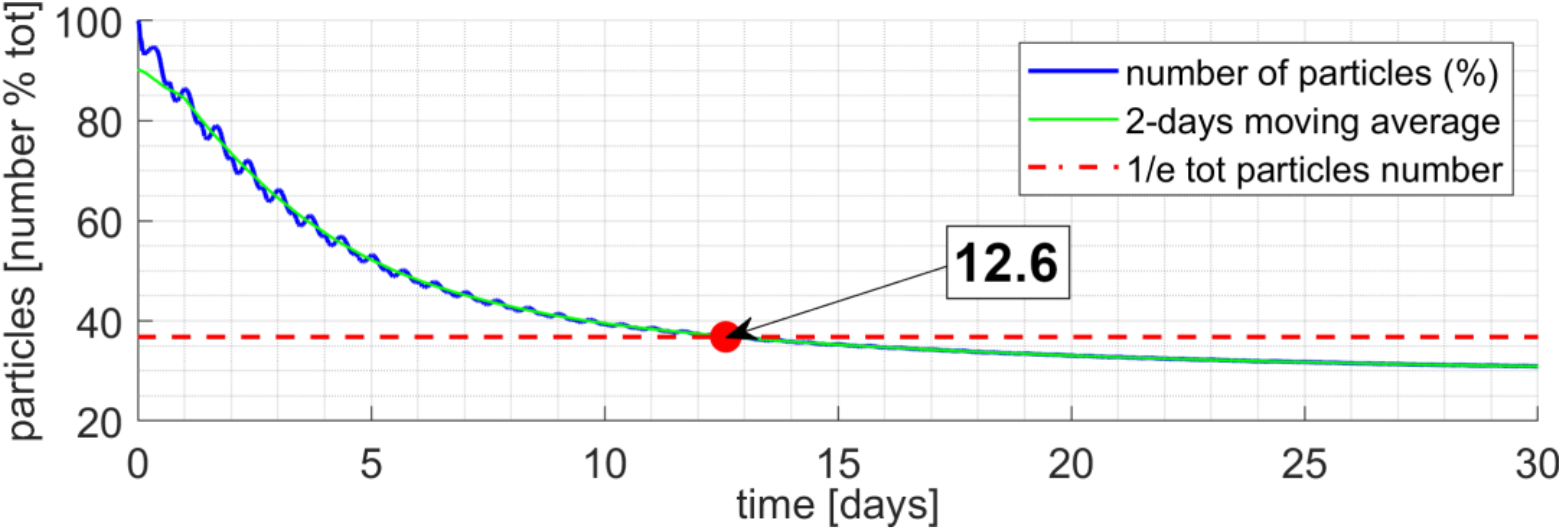
The plot shows the number of particles located in the estuary divided by their initial number as a function of time. The red circle indicates the time at which the number of particles reaches 1/e of the initial value. That time is our estimate of the estuary-scale residence time *(RT_E_*). The white box contains the value of the residence time expressed in days.

### 3.5. Filtration Services

Figure 5A shows the distribution of the *FS* calculated for the oyster reefs in the estuary *(FS_R_*). The boxes in Figure 5A show the distribution of the *FS* calculated at the watershed scale *(FS_W_).* The proportion of the estuary cleared by a single reef *(FS_R_*) varied from 0 to 0.90% across the estuary. The total volume of the estuary cleaned by the oyster reefs over a estuary residence time *(FS_E_*) was ~60%. The greatest contribution to the filtration of the GTM estuary was provided by watershed W1 (~20% of *FS_E_*). A large contribution to estuary biofiltration was provided by reefs in watersheds W6, W7, and W8, providing 6.07%, 10.12%, and 9.23% of the *FS_E_*, respectively. Lower *FS_E_* values for the watersheds were obtained from W3, W5, and W9, which provided 1.49%, 2.46%, and 4.16%, respectively. The lower values were likely due to their more peripheral location of the watersheds, away from the major channels connecting the inlets. The lowest value (0.28%) for watershed W4, both because W4 had high tidal velocities, generated by the water exchange with the ocean through the St. Augustine inlet, and because it contained a relatively small number of reefs. In addition, it is important to notice that, at the end of the simulated estuarine residence time, ~22% of the mass initially contained in the estuary, left the GTM from the inlets of St. Augustine and Matanzas, as well as from the southern and northern boundaries of the ICW. For this reason, the total exchange of material of the estuary corresponds to ~81% per tidal cycle.

**Figure 5.**
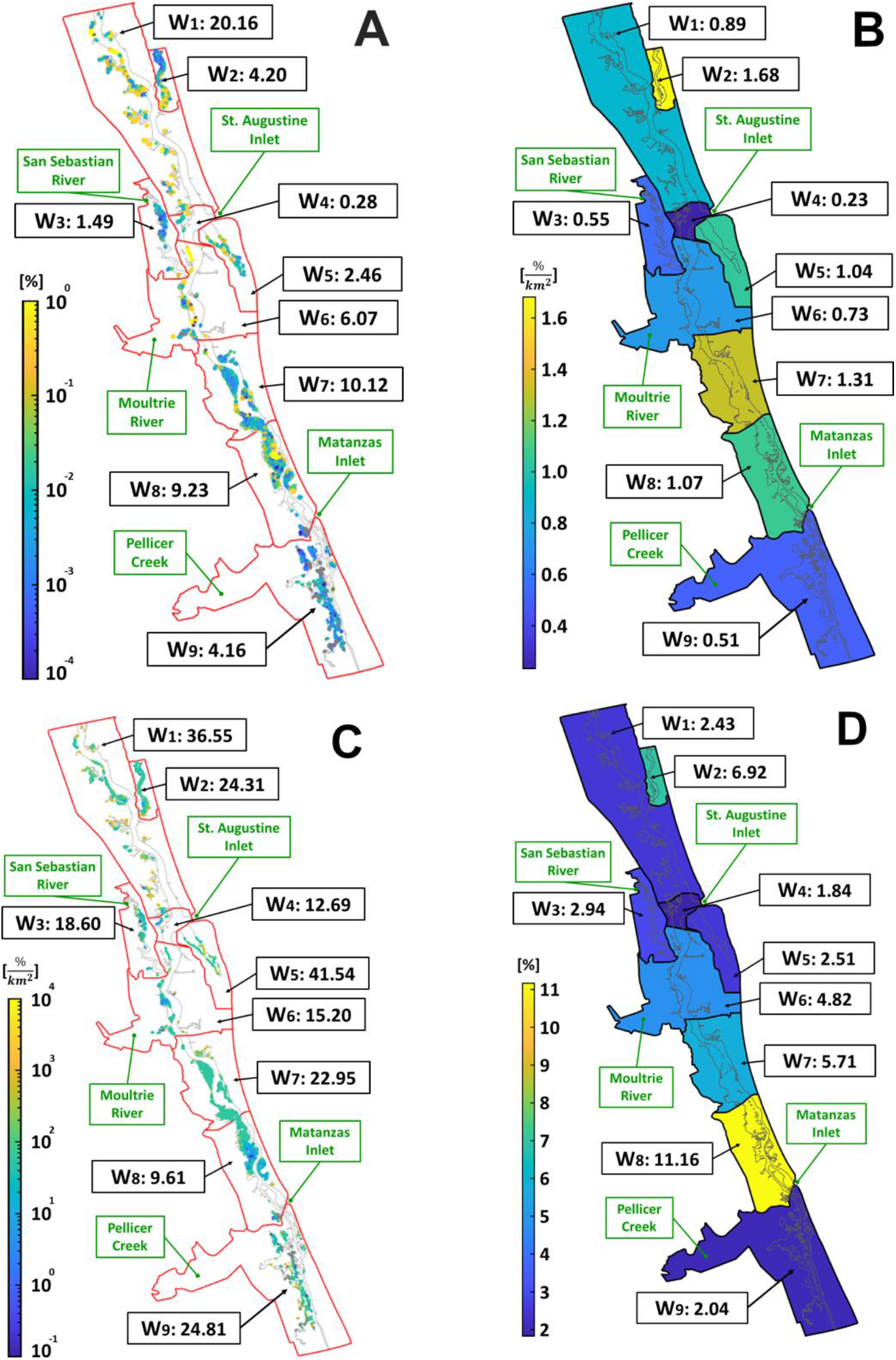
(A) Filtration Services at the reef scales *(FS_R_*) and the watershed scale *(FS_W_*) for each watershed (Wi, i=1,9). Both *FS* are reported in percentage of estuary filtered within a residence time [%]. (B) The spatial distribution of *FS_W_* per square kilometer of intertidal watershed area 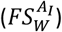. For each watershed (Wi, i=1,9), the values are reported in percentage per square kilometer of intertidal area of the watershed [%/km^2^]. (C) The spatial distribution of the Filtration Services at the reef scale per unit of reef area 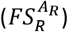, and its average values per each watershed 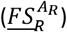. For each watershed (Wi, i=1,9), the values are reported in percentage per square kilometer of reef area [%/km^2^]. (D) The spatial distribution of the percentage of intertidal watershed area occupied by oyster reefs 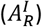. The values are reported in %.

Although Figure 5A provides the total *FS* for each watershed, it had two drawbacks. The first drawback was that for spatially uniform oyster reef density and hydrodynamics, *FS_W_* increased with the watershed area *(A_I_*), and in particular with the intertidal watershed area, where oyster reefs preferentially develop. Therefore, Figure 5A does not allow us to understand if large watersheds provided a large service because of their size or because of the filtration capability of their reefs. To overcome this issue, we calculated 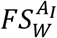 (Figure 5B), which was the *FS_W_* per unit of intertidal watershed area *A_WI_*. The latter is shown in Figure 2C. The second drawback was that the filtration service at the reef scale *FS_R_* increased with the reef size. Analogously to *FS_W_,* it was not clear if a large *FS_R_* in Figure 5A indicated a specific ability of the reef to filter water, or it was a consequence of a large reef size. To estimate the relative contribution of each reef to *FS_W_* and *FS_E_* independently from their size, we calculated 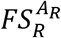, which were the values of *FS_R_* per unit of reef area *A_R_*. Figure 5C reports the spatial distribution of 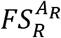 in the GTM and its watershed-averaged value 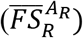, computed by using the reef areas as weights:

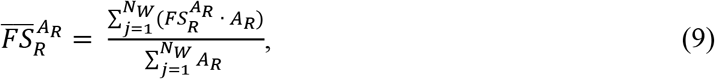

where *j* spans over the *N_w_* reefs in the watershed *W.* Finally, in order to estimate the propensity of reefs to establish in each watershed, we calculated the area of oyster reefs per unit of intertidal watershed 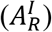.

Computing the values of the various *FS* per unit area allowed us to (i) compare the relative contribution of each reef and watershed to *FS_E_*, and (ii) identify which region of the estuary could provide the maximum increase in *FS_E_* if targeted for restoration. In short, *FS* per unit area described the filtering efficiency of reefs and watersheds. *FS* at the watershed scale (Figure 5A) can be written as:

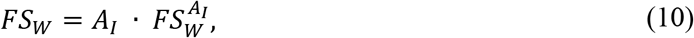

Therefore, the *FS* at the watershed scale increased with *A_WI_* and 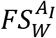. Figures 5A and 2C show that *FS_W_* mainly increases with *A_I_*, thus partially hiding the effect of 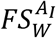. Exceptions were watersheds among those with extremely similar intertidal areas, such as W6 with W7, W8 and W9. *A_I_* was larger for watershed W6, W8 and W9 than for watershed W7 (8.283, 8.596 and 8.239 vs 7.720 km^2^). However, W7 had a greater 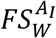, due to the greater number of reefs located in it, and their larger individual contribution to *FS_E_*. Other exceptions were W2, W3 and W5. *A_I_*, was larger for watershed W3, followed by W2 and W5 (2.728, 2.496 and 2.357 km^2^, respectively). However, 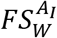 is higher for W2, followed by W5 and W3. Moreover, the small values of *FS_W_* in Figures 5A for W2 and W5 may be misleading. These small values were not due to poor filtration capability of the reefs but rather their small area. In fact, W2 and W5 displayed the first and fourth highest value of 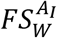 (Figure 5B).

We can further break down *FS_W_* to mechanistically understand the drivers of the observed values. Since by definition:

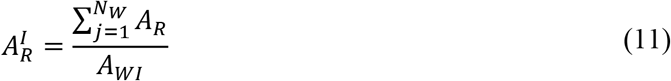

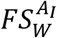 reads:

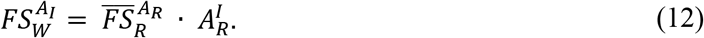

Therefore, the watershed scale *FS* per unit of watershed wetted area (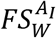, Figure 5B) increased with the average *FS* of the reef per unit reef 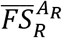(Figure 5C) and area of oyster reefs per unit of wetted watershed 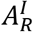 (Figure 5D). Finally, by substituting (12) in (10), we have that:

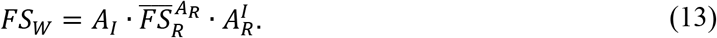

Which provided the full dependence of *FS_W_* (Figure 5A) on the three quantities shown in Figure 2C, 5C, and 5D.

Figures 5B, C, and D show that large values of 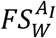, were mostly associated with large 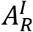. The three greatest 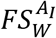 in the estuary, were obtained from watershed W2, W7 and W8, were associated with the three greatest 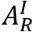 in the GTM, equal to 6.92%, 5.71% and 11.16%, respectively. The corresponding 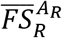 values for W2, W7 and W8 were 24.31, 22.95 and 12.69 %/km^2^, respectively. Notably, although W8 had the lowest observed value of 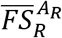, the 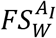 value was the second greatest due to a large density of reefs in the watershed. The lowest 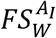 in the estuary, obtained for watershed W4, was associated with the lowest 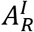(1.84%), and a second lowest 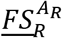 (12.69 %/km^2^). A similar 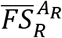, equal to 15.20 %/km^2^ was observed for watershed W6. However, due to a higher presence of reefs in W6 than in W4 (4.82% vs. 1.84%), a greater 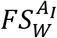 was observed for W6. For watershed W1, W5, and W9, the 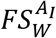 was associated to the greatest values of 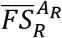 observed in the estuary, equal to 36.55, 41.54 and 24.81 %/km^2^, respectively. However, the contribution of these watersheds to 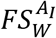 was reduced by the low density of reefs 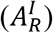, which was equal to 2.43%, 2.51%, and 2.04%, respectively. The moderate values of 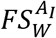 observed for watershed W3 was due to the combined effect of intermediate values of 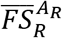 and 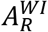.

Figure 5C shows how the contribution of the singular reef to 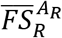 varied in the estuary. In particular, the lowest 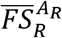 were associated with watersheds close to the inlets (W4 and W8), and with watersheds mostly covered by the main rivers of the GTM (W6). Conversely, the greatest 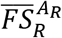 were associated with watersheds comprising the ICW lateral branches (W2, W3, and W5), or mostly covered by salt marshes (W1, W7, and W9).

### 3.6. Genetic algorithm

A relationship to describe 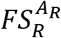 could be obtained for the *j^th^* reef by substituting Equation (5) in (7). After some steps we obtain:

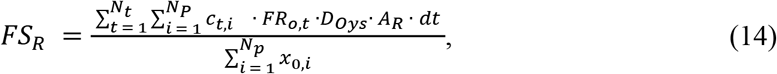

which could be written as:

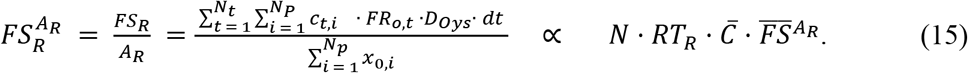

The relationship in Equation (15) shows that 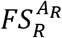 depends on the predictors described in Section 2.6.1, which account for the main properties of the reefs, and of the particles entering them.

Table 2 shows the value of the statistical parameters obtained for each predictor in Equation (15), when they were used individually as predictor in a linear regression describing 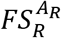, assuming that the other three parameters in (15) are constant. The statistical parameters show that the effects of 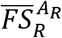, *RT_R_*, and *N* on the reef-scale 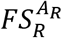 were negligible. Notice that, for 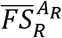 the calculated p-value was lower than 0.005, indicating statistical significance; however, the low R^2^ and the high RMSE and MAE, confirmed the negligible contribution of this predictor. For *N* and *RT_R_*, both p- values were greater than 0.05, and the almost null R^2^ confirm their negligible contribution to identify a relationship describing 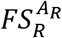. The best predictor of 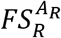 was 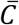 as it was both highly significant (p-value < 0.005) and explained more than one fourth of the variability (R^2^ = 0.257). This relationship suggested that the *FS* were influenced by downstream effects in the estuary, because <u>**C**</u> accounts for the filtration history of the particles. This was confirmed by the value of RMSE, which was the lowest calculated among single predictors models (72.82). Additionally, the value of MAE, was comparable to the ones obtained for the other predictors (29.41 vs. 34.05, 33.96, and 34.77), indicating that the relationship obtained from *<u>C</u>* to describe 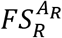 reduce the variance of the distribution of error magnitudes, but not the average magnitude of the errors.

**Table 2.** For each single predictor are reported the p-value, R^2^, Root Mean Square Error (RMSE), and Mean Average Error (MAE) calculated from the relationships obtained from the Genetic Algorithm to describe 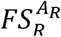. The relationships are obtained by using each predictor described in Section 2.6.1, individually. A full, multiplicative model was obtained by considering most predictors described in Section 2.6.1, in the GA.

| Predictor | p-value | $R^2$ | RMSE | MAE |
| --- | --- | --- | --- | --- |
| $\overline{FS}_{Ar}^{Ar}$ | <0.005 | 0.0024 | 84.36 | 34.05 |
| $RT$ | 0.878 | 0.0047 | 84.26 | 33.96 |
| $\bar{C}$ | <0.005 | 0.257 | 72.82 | 31.53 |
| $N$ | 0.249 | 0.0013 | 86.20 | 34.77 |
| $N \times \bar{C} \times \overline{FS}_{Ar}^{Ar}$ | <0.005 | 0.897 | 27.97 | 12.11 |

When the model predictors in Equation (15) were combined in the GA to determine a relationship describing 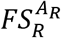, the GA suggested the following relationship:

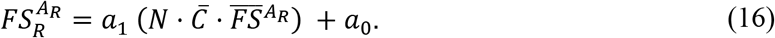

Because 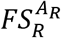 was null when the number of particles entering the reefs *(N),* and their average concentration 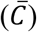 are zero, as well as when the filtration capability of a reef per unit of reef area 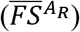 was null, we performed the regression setting *a*_0_= 0. We obtained a value of *a*_1_ equal to 7.1535. Table 2 shows the value of the statistical parameters obtained for this relationship. This full model was both highly significant (p-value <0.005) and had much greater explanatory power (R^2^ = 0.897) than any single-parameter model (Figure 6). The RMSE and the MAE decreased to 27.97 and 12.11 respectively, showing a strong reduction of the prediction error.

**Figure 6.**
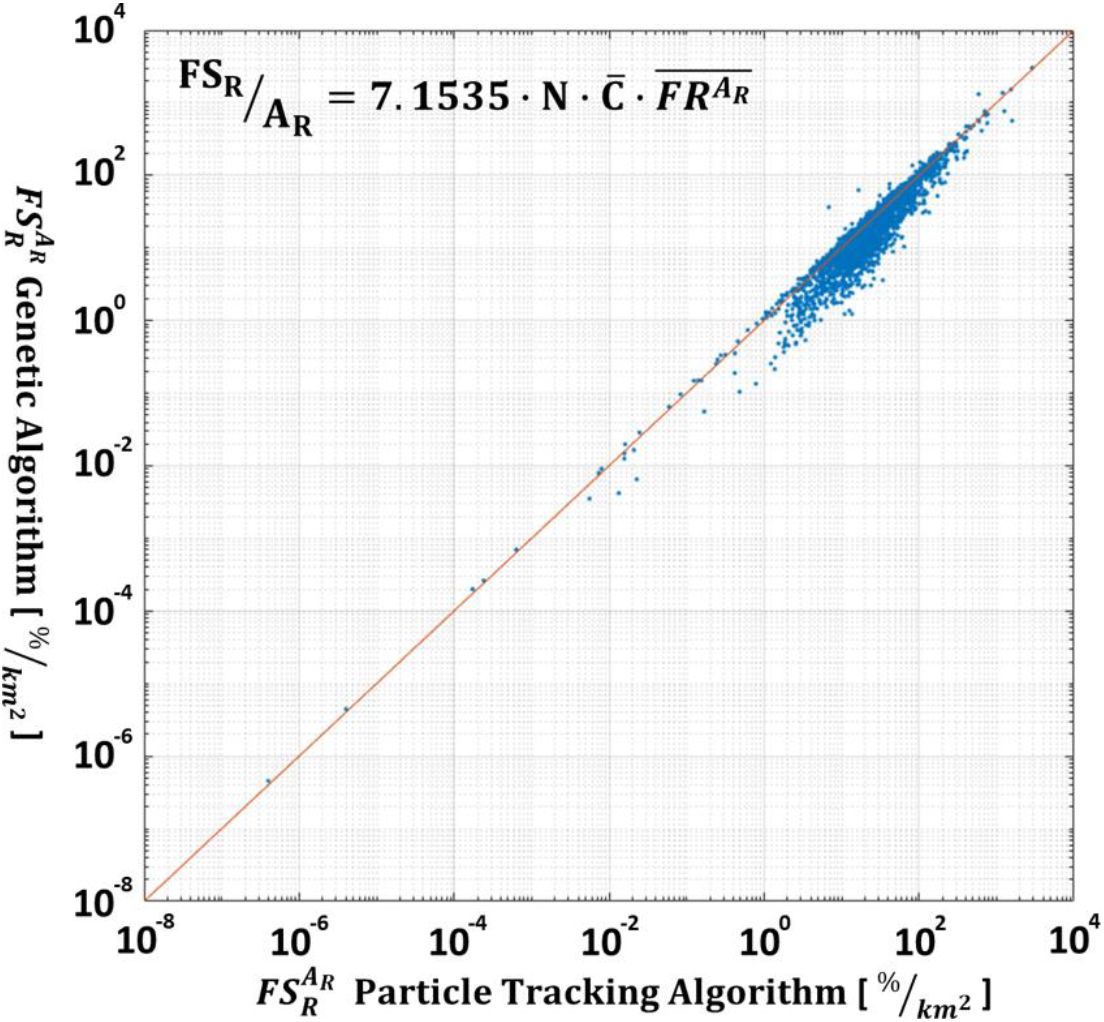
Scatter plot comparing the values of the FSRAR obtained from the MATLAB Algorithm based on the particle tracking model (x-axis) and the FSRAR obtained from the relationship obtained from the Genetic Algorithm (y-axis). The relationship is reported in the figure. Both axes are on a logarithmic scale to enhance the visibility of the point cloud.

The relationship obtained from the GA suggests that the average initial concentration (*C*) of the particles entering a reef, as well as their number (*N*), were more important than their permanence over the reef, which value corresponds to the *RT_R_*. The limited importance of *RT_R_* was due to its limited variability observed over the reefs. In fact, ~80% of the reefs showed an *RT_R_* between 1 and 5 minutes (Figure 7). Similarly, the average concentration had a greater impact than the average mass of the particles entering a reef. This was because 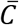 accounted for the dilution of the particulate in the seawater volume above the reefs, which highly influenced the *FS* of the reef. The good prediction capabilities of 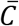 were confirmed by the statistical parameters reported in Table 2. Moreover, we wish to highlight that the value of 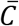 depends on: (i) the number of reefs crossed by the particles throughout the estuary. This number in turn depended on local- and estuary-scale hydrodynamics, initial distribution of particles injected in the estuary, and spatial distribution of estuary reefs; (ii) the local reef filtration rate per unit of area 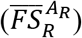, which in turn depended on the local oyster population characteristics. Thus, by considering 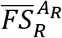, Equation (16) underpinned the importance of the oyster reef properties in the resulting *FS*. Finally, although the model was initiated with uniformly distributed particles over the GTM, downstream effects were not uniformly distributed. For this reason, in the GTM, the oyster reef contribution to water quality depended on their spatial arrangement and upon the estuary hydrodynamics.

**Figure 7.**
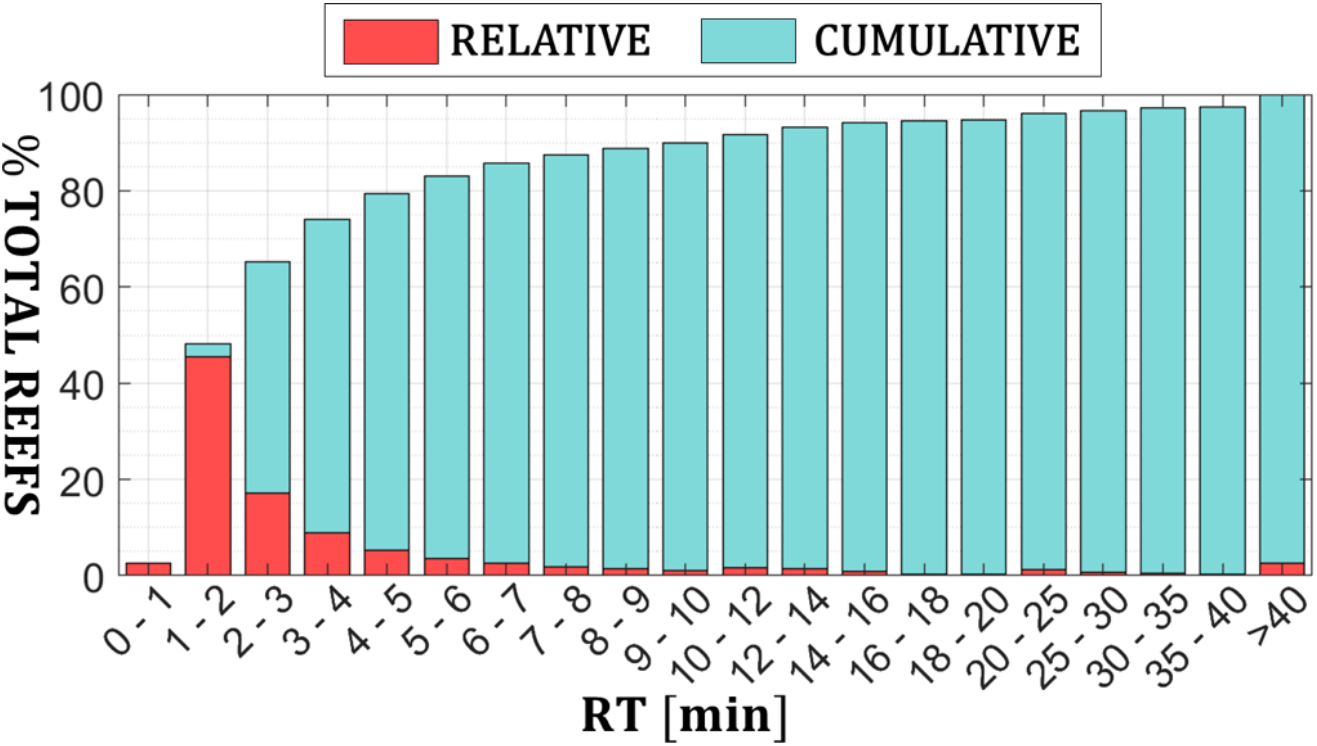
Relative (red) and cumulative (teal) frequency distribution of the residence time calculated for the reefs in the estuary.

## 4. Discussion

The native oyster population in GTM is exceptionally intact and robust when compared to many other populations in the US and elsewhere that are either in poor condition or functionally extinct (Beck et al. 2011). Importantly, our ability to describe this population and estimate the ecosystem services conferred by this population were bolstered by detailed surveys of 240 randomly selected reefs of the approximately 4,300 in this system, which supplied demographic and density information across all watersheds and subestuaries. Furthermore, because we could resolve how services varied among populations after controlling for reef size, we could both determine which populations were most efficient at filtering particulate and tease apart the role of various hydrodynamic factors governing filtration services. These types of analyses can inform future management of these populations and preservation of their valuable services.

For comparative purposes, we specifically chose to estimate the clearance rates of *C. virginica* by following the approach of zu Ermgassen et al. (2013), who modeled the current and historic filtration services of this species throughout the Atlantic and Gulf Coasts of the US. Although our approach to modeling the physical interactions between populations and suspended particles differs significantly between these studies, comparing results of these studies helps illustrate how the filtration services estimated for present-day GTM populations surpassed those of all contemporary and historic populations estimated by zu Ermgassen et al. (2012). Indeed, among the 13 estuaries modeled, the maximum filtration services were estimated to be contributed by historic populations in Matagorda Bay during the Fall (51% bay filtered within a residence time). All other peak filtration services estimates among the other historic populations were much less apparent (mean: 4.5%) and present day services among these same populations were on average a small fraction of the historic services (−71% of historic value). The impressive filtration services of oysters in the GTM estuary, along with relatively short residence times (Phlips et al. 2004), likely play a major role in keeping phytoplankton biomass low and providing resilience to natural and human disturbances (N. Dix et al. 2013).

The estimated population metrics and reef-scale biofiltration rates underpinning the ecosystem scale results are also worth examining. Reefs were dominated by relatively small individuals (mean shell height = 35.02 mm) due to persistent annual recruitment; however, oyster densities within reefs were also quite high (mean = 1855 ind./m^2^) and relative abundance of these populations within subestuaries (mean = 52 ind./m^2^ of estuary) was relatively high compared to historic coverage across the 13 estuaries (mean historic oyster coverage: 36.6 ind./m^2^) examined by zu Ermgassen et al. (2012).

It is important to note that we only account for oyster filtration services within this model while neglecting those reef community members that also contribute to filtration services. Indeed, other GTMNERR survey data indicated a strong relationship between oysters and other suspension feeders including ribbed mussels (R^2^ = 0.69) and regular presence of other filter-feeding invertebrates (i.e. quahog clams, barnacles, mahogany date mussels) (Marcum et al. 2018). Non-oyster suspension feeders on reefs can add appreciably to total reef biomass (e.g. ~16%) and contribute significantly to biofiltration and water quality improvement (Kellogg et al. 2013). That said, the relatively average filtration rates observed (mean 2.54 l h^-1^) combined with the high density of oysters on reefs produced reef-scale filtration rates (mean 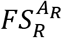: 4,136 1 h^-1^ m^2^) that were orders of magnitude greater than maximum filtration rates indirectly measured on natural (44 1 h^-1^ m^2^) and constructed (154 1/h/m^2^) reefs in the Gulf of Mexico; albeit these reefs were much less dense (407 ind. m^-2^ and 690 ind. m^-2^, respectively) than those found in GTM (Milbrandt et al. 2015). Nevertheless, we are confident that if other community members were included, it would not be surprising to observe our reef-scale filtration estimates increase substantially. The large estimates for 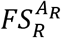 in GTM are not without precedent and resemble the maximum (summer = 26°C) filtration rates estimated for pre-colonial reefs in the Chesapeake Bay with a maximum age between 14 y and (3,872 1 h^-1^ m^2^) and 16 years old (5,388 1 h^-1^ m^2^) (Mann et al. 2009). Maximum filtration rates are appropriate to use in this comparison as the average temperature in each subestuary of the GTM in the model (~24.5°C) approached that which elicits maximal feeding responses of *C. virginica* (e.g. 26°C Newell et al. 2005), 27°C zu Ermgassen 2013/ Cerco and Noel 2005).

In many previous models of oyster filtration services, the density, demographics and precise spatial distribution of reefs (historic or otherwise) are unknown and many assumptions about the access populations have to overlying water must be simplified during model creation (e.g., Pomeroy et al. 2006; Fulford et al. 2010; zu Ermgassen et al. 2013; zu Ermgassen et al. 2013). Our approach overcame this limitation by using a coupled FWRI+GTMNERR dataset, which contained detailed and up-to-date spatial and biological information of the reefs in the study area. However, while the FWRI dataset contains the spatial location and the extension of all the known reefs occupied by living oyster communities in the GTM estuary, the biological information in the GTMNERR dataset of area were available only for a limited number of reefs (~6% of total reefs). We verified the accuracy of the IDW interpolation method used here to estimate the biological properties of the oysters in unsurveyed reefs, by excluding 20 reefs and using them to compute the error. MAE and RMSE were equal to 6.3 and 6.1 mm for the shell height, and to 578.6 and 310.3 oysters/m^2^. This was due to a fairly uniform distribution of the surveys, which spanned the whole GTM estuary and its tributaries. Future work will include examining how within reef variability, whether due to natural processes or harvest practices on demographic properties, alters the estimates of *FS* produced by reefs and populations.

Because of the high computational cost of numerically tracking particles over each reef in large domains, researchers developed simplified methods to evaluate oyster *FS* based on coarse regular grids (e.g. Gray et al. 2019). The use of these grids has two major drawbacks: (i) the boundaries of a naturally-irregular reef morphology cannot adequately be described by a coarse regular grid; (ii) particles entering a coarse cell can be filtered even if they do not directly travel over the reef (Figure A1). To overcome these drawbacks, our approach modeled the hydrodynamic in the complex GTM estuary using a high-resolution curvilinear grid, which follows the main watercourses, coupled with high-resolution elevation data, obtained from open datasets and targeted local surveys. More importantly, our model filtered only the particles traveling over the reefs in the GTM estuary. The small increase in the computational costs related to this method was completely justified by the improvement in the description of the filtration history of the tracked particles. By identifying a clear correlation between the concentration of water parcels traveling over an oyster reef and the reef *FS,* we showed that downstream effects directly influenced *FS*, and have to be explicitly considered when planning restoration if water quality improvement is a major project goal. Restoring oysters by prioritizing locations with relatively high residence time as in Gray et al. (2019) might not always be the exclusive best strategy. Our study indicated the optimal locations for targeted restoration were determined by taking into account both residence time and water refiltration through downstream effects, which can only be accomplished with precise spatial knowledge of oyster locations, abundance, and hydrological patterns. Note that we used uniformly distributed particles over the GTM to make the results obtained for each reef in the estuary independently of the initial position of the particles. However, the reef- and the watershed-scale *FS* calculated for the GTM had a non-uniform distribution due to the complex local and estuary-scale hydrodynamics. A high-resolution modeling approach was then needed to precisely describe the GTM hydrodynamics, and to correctly estimate the contribution of oyster reefs to estuarine water quality.

### 4.1. Future developments

We estimated the local contribution of individual reefs to the global (i.e. estuary-scale) *FS*. This is a fundamental step forward for planning ecological conservation actions in the GTM estuary and represents an approach that others from outside the GTM may wish to adopt when selecting locations for reef creation, enhancement, or conservation. Future applications of our model include (i) evaluating the impact reef filtration has on pollutant sources in the GTM, (ii) understanding how harvesting oysters shifts population demographics (including within-reef density and size frequency distributions) and impacts subestuary- and estuary-scale *FS*, and (iii) describing the growth of the reefs with population dynamic models, and consequently their short- and long-term survivability under different management, biological, and hydrodynamic scenarios. We will apply the model using point and nonpoint pollutant sources. Simulations will include point sources such as wastewater plant discharge, or distributed sources, such as septic tanks or agricultural runoff. The release could be either due to a programmed operation, or accidental, if due to an unattended leakage or as a consequence of extreme weather events (such as hurricanes and Nor’easters). Alternatively, if a population is damaged or overharvested, this model could be useful to help understand larval supply and transport which may be leveraged to restore and repair reefs. Gray et al. (2019) showed that prioritizing oyster restoration in regions with large residence time and high encounter rates that promote refiltration can help resource managers achieve ecological restoration goals with less resource investment than deploying oyster randomly within the habitat. We agree that accounting for hydrodynamics can improve ecological outcomes and resource use efficiency during oyster restoration; however, our genetic algorithm showed that the average mass concentration of the particles entering the reefs *(C_in_),* representing downstream effects, was better than residence time alone at estimating *FS* at the reef scale in a given location. Therefore, more research is needed to develop a reliable approach that maximizes *FS* by taking into account not only residence time but also downstream effects.

## Conclusions

In this work, we used a numerical model that solved hydrodynamics and transport of particulate matter to estimate oyster *FS* of Eastern oysters (*C. virginica*) in the GTM estuary, FL, which possess traits (reef density, oyster abundance, etc.) that may resemble “pristine” populations that were more common in the US prior to arrival of Euro-American settlers. The output of the numerical model consists of the particles’ location at each time step of the simulation. This output is read by a Matlab script, which tracks the average time the particles spend over the cells of a regular 50m×50m grid, the watersheds, and the estuary. We used this information to estimate the residence time and the *FS* at different scales (local, watershed, and estuary). For this purpose, results show that the local- (i.e. at the 50m×50m scale, *RT_L_*) and watershed-scale (*RT_W_*) residence times were not uniformly distributed in the GTM. *RT_L_* was minimum at locations close to inlets (~0-2 minutes), increased through the rivers (~1-7 minutes), and reached its maximum value in the marshes (~7-240 minutes). At the boundary between the intertidal and subtidal areas, where the oyster reefs are mostly located, *RT* ranges between 1 and 5 minutes. *RT_W_* depended on the dimension of the watersheds, their proximity to the inlets, and the area of the watershed occupied by salt marshes. The estuary residence time was ~13 days. By tracking the time spent by each particle over a reef, the model accounted for the mass removed from the particles floating over the reefs. Accounting for reef area when estimating *F S* provided novel insight of the relative contribution of reefs which can provide valuable resource management information. The model results show that: (i) oyster reefs populating the GTM improved water quality by filtering ~60% of the estuary’s volume within a single residence time; (ii) the spatial distribution of the filtration service at the reef and watershed scales varied spatially across estuary; (iii) at the watershed scale, *FS* depended on the distribution of the reefs in the watershed, and on the proportion of the wetted watershed area they occupy. Finally, we used a Genetic Algorithm to identify the predictors that best described the reef-scale filtration rates. Our genetic algorithm revealed that the average mass concentration of the particles entering the reefs *(C_in_*, a proxy for downstream effects), rather than residence time, best described the reef-scale contribution to estuary-scal *FS*. In future research projects, we intend to apply the model in a variety of ways to explore how natural and anthropogenic effects influenc *FS*.

## Acknowledgements

This research was partially funded by NSF BIO-OCE Award 1736943 to DLK

## Supplementary Material

### Appendix A

#### Numerical simulation period

To compute the average residence time of the estuary, we simulated 30 days in 2018. The simulation period goes from May 9^th^, 2018 to June 10^th^, 2018. We choose that period because it contains the most representative spring and neap tides of the year. We use the following procedure to determine the simulation period: (i) we reconstruct the astronomic signal for 2018 using the harmonic tidal components of the local NOAA station “St. Augustine Beach, FL” #8720587”. We choose this station because it is the only one contained in the study domain, (ii) We calculate the tidal ranges in 2018 using consecutive low and high tide levels extrapolated from the astronomic tidal signal. (iii) We classify the tidal ranges using the 25^th^ and 75^th^ quantiles of their distribution: we consider that the ranges lower than the 25^th^ quartile belong to neap tides, and the ranges greater than the 75^th^ quantile belong to spring tides. We then calculate the average tidal range for neap and spring tides. (iv) We then divide the 2018 astronomic tide into groups of 30 consecutive days, containing two spring and two neap tides. (v) For each group, we identify the tidal ranges associated with spring and neap tides by using the quantiles we previously identified for 2018. Then, for each group, we calculate the average tidal range for neap and spring tides, the difference between these average values and the ones calculated for 2018, and the sum of these two differences. The group with the lower value of this sum contains the most representative couple of spring and neap tides of 2018.

#### Harmonic constituents

We collected the tidal harmonic constituents from the NOAA stations “St. Augustine Beach, FL, #8720587”, “Jacksonville Beach, FL, #8720291”, and “Daytona Beach (Ocean), #8721020”. Jacksonville Beach station is placed 17 km north of the northern boundary of our study domain. Dayton Beach station is placed 45 km south of the southern boundary of the study domain. St. Augustine station is the only one located in the study domain, and it is placed 6 km south of the homonymous Inlet. All the stations are placed close to the coastline. The considered harmonic constituents are M2, S2, N2, K1, M4, O1, M6, NU2, MU2, M1, J1, SSA, SA, Q1, T2, R2, P1, L2, and K2. We applied the harmonics relative to St. Augustine station on the offshore boundary at the same latitude as the station (central red square in Figure 1A). We determine the harmonics at the northern and southern limits of the offshore boundary (red squares in Figure 1A) by linearly interpolating the harmonics at St. Augustine with the harmonics at Jacksonville and Daytona Beach, respectively. Finally, at the northern and southern cross-shore boundaries of the numerical model, we applied a Neumann boundary condition following the method proposed by (Roelvink and Walstra 2005)

